# Anatomical correlates of face patches in macaque inferotemporal cortex

**DOI:** 10.1101/2020.06.10.144600

**Authors:** Michael J. Arcaro, Theodora Mautz, Margaret S. Livingstone

**Affiliations:** Department of Psychology, University of Pennsylvania; Department of Neurobiology, Harvard Medical School

## Abstract

Primate brains typically have regions within the ventral visual stream that are selectively responsive to faces. These face patches are located in similar parts of macaque inferotemporal (IT) cortex across individuals though correspondence with particular anatomical features has not been previously reported. Here, using high resolution functional and anatomical imaging, we show that small “bumps” along the lower bank of the superior temporal sulcus are predictive of the location of face-selective regions. Recordings from implanted multi-electrode arrays verified that these bumps contain face-selective neurons. These bumps were present in monkeys raised without seeing faces and that lack face patches, indicating that these anatomical landmarks are predictive, of but not sufficient for, the presence of face selectivity. These bumps are found across primate species, indicating common evolutionary developmental mechanisms.

## Introduction

Prior research has demonstrated a relationship between cortical folding and the functional organization of primary sensory areas (Kaas et al., 1979; Fischl et al., 2008; Rajimehr and Tootell, 2009; Da Costa et al., 2011). Within the visual system, the calcarine sulcus not only serves as a macroanatomical landmark for primary visual area V1 (Holmes, 1918; Hinds et al., 2008), but the folding patterns within the sulcus are predictive of the retinotopic organization (Schira et al., 2009; Benson et al., 2012). Given the complexity and variability in cortical size and shape across individuals, traditionally, it has been thought that there is little correspondence between cortical folding and visual areas beyond V1. However, research over the past decade has revealed a surprising degree of structure-function correspondence across the cortical surface (Amiez and Petrides, 2014; Benson et al., 2014; Weiner et al., 2014; Witthoft et al., 2014; Leroy et al., 2015).

Primates typically develop several regions within inferotemporal cortex (IT) that are selectively responsive to faces. Three spatially distinct sets of face patches, the posterior lateral (PL), middle lateral (ML), and anterior lateral (AL), have been identified along the lower bank of the superior temporal sulcus (STS) in macaques (Tsao et al., 2006; Tsao et al., 2008). Two other patches middle fundal (MF) and anterior fundal (AF) lie further down the sulcus at the same AP location as ML and AL, respectively. These 5 face patches are located in similar parts of IT across individuals, though correspondence with particular anatomical features has not been observed.

Here, we performed high-resolution anatomical neuroimaging on eighteen rhesus macaques. Seven monkeys were raised with normal visual experience of faces and developed face patches identified by fMRI within the first year of life. A topological analysis of each monkey’s cortical surface revealed that face patches PL, ML and AL were localized to focal convex protrusions along the STS, which we refer to as bumps. Neural recordings from five monkeys confirmed that each bump contains face-selective neurons and demonstrate that targeted recordings of face-selective neurons in the macaque brain can be achieved with high success without requiring fMRI. These bumps were morphologically similar in a separate group of monkeys raised with an abnormal visual experience of faces and that lacked face patches. Thus anatomical landmarks may predict the location of functional specializations but are not sufficient for their presence. Using publicly available datasets, we identified these bumps *in utero* in rhesus macaques as well as postnatally in several other primate species. This suggests that bump formation emerges from general mechanisms ubiquitous across the primate order. These general mechanisms may underlie the organization of maps, areas, and patterns of architectonics, which in turn may influence functional specializations.

## Results

To probe the relationship between face selectivity and the anatomical topology of inferotemporal cortex (IT), we performed functional and anatomical MRI on seven macaque monkeys reared with normal visual experience. Functional scans were aligned to high resolution (0.5mm isotropic) T1 anatomical MR images. Regions preferentially active to images of faces vs. objects were identified along the lower bank of the superior temporal sulcus (STS) in the right (Figure 1) and left (Figure 1 – figure supplement 1) hemispheres of each monkey. Face-selective responses appeared to co-vary with the topology of the STS. The lower bank of the STS is not a uniformly smooth sulcus, rather there are several focal protrusions along the posterior-to-anterior axis, which we refer to as bumps, where the cortical surface bulges. These bumps are most apparent in parasagittal sections of the STS. When viewed coronally, the bumps are less prominent relative to the broader convexity of the temporal gyrus. These bumps were visible on both high resolution T1 scans and lower resolution functional EPI images (Figure 1 – figure supplements 2 & 3). Across monkeys, face selectivity consistently fell on three bumps along the posterior-to-anterior extent of the STS, which we refer to as the posterior (Figure 1, green line), middle (pink line), and anterior (blue line) bumps. More broadly, there was no consistent relationship between surface curvature and the magnitude of the faces vs. objects contrast along the entire STS (t(13)=2.0268, *p* = 0.0637). The lack of a positive relationship is likely due to the presence of additional bumps along the STS that are not face selective as well as the prominent mediolateral convexity of the temporal gyrus. Similarly, neither cortical thickness nor sulcal depth were significantly related to face selectivity within the STS (ts(13) < 0.78, *ps* > 0.45). Together, this indicates that face selectivity was specific to these three anatomical landmarks and does not reflect a broader relationship between convex cortical folding and face selectivity.

**Figure 1.**
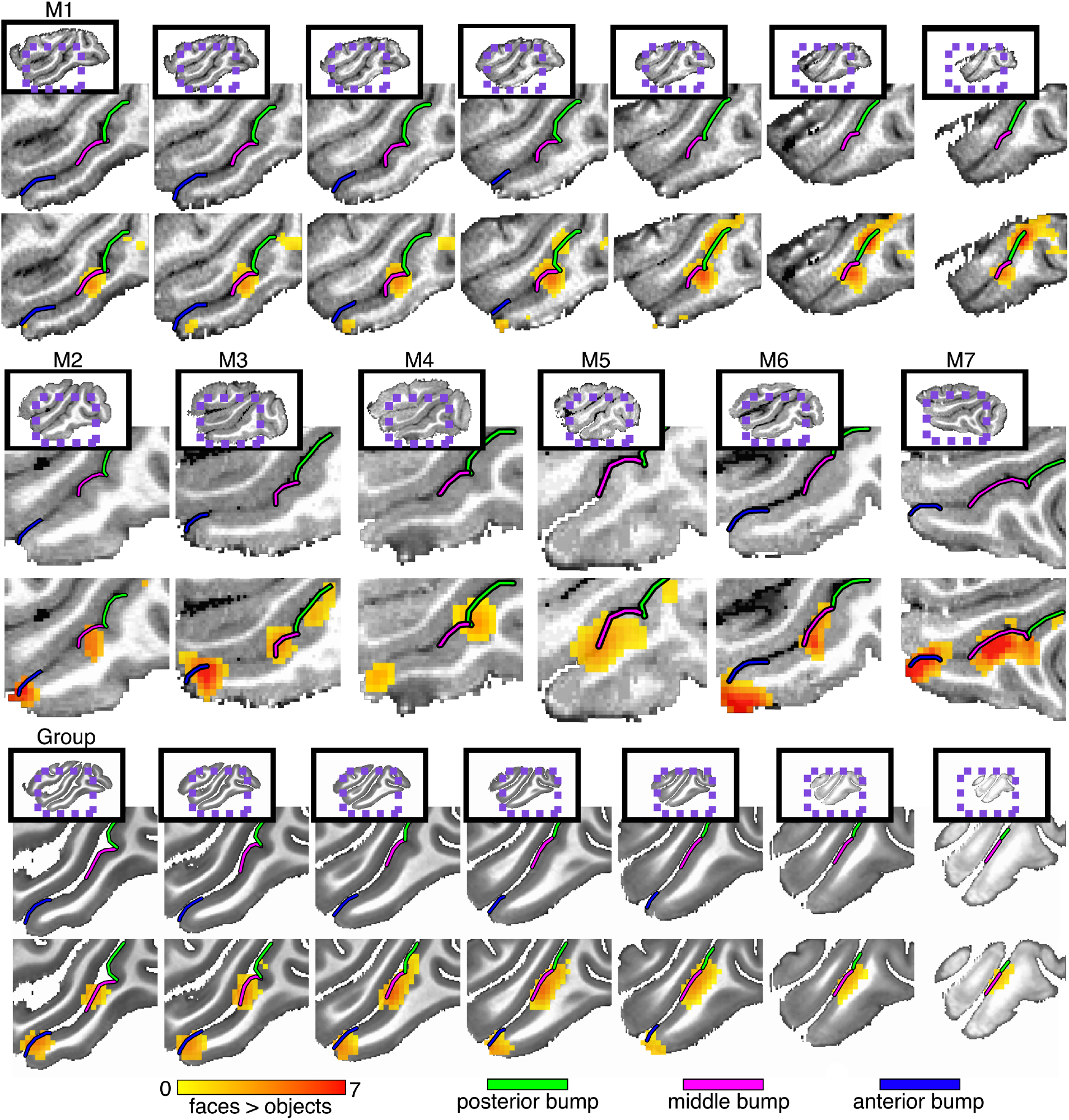
Anatomical localization of face selectivity in right hemisphere STS. Preferential activity for faces vs. objects was identified along the lower bank of the STS in all 7 monkeys reared with normal face experience (*p* < 0.0001, FDR-corrected). (top) Seven sequential sagittal slices at 1mm spacing in the right hemisphere of Monkey 1. (middle) Single sagittal slices showing localization of face-selective activity to bumps in the right hemispheres of the other normally reared monkeys. (bottom) Group average face selectivity falls on anatomical bumps in the STS. Seven sagittal slices at 1mm spacing in the right hemispheres of the NMT template. Group average faces vs. objects (threshold to show only surface nodes where at least 3 monkeys showed significant activity) and convexity maps on cortical surface reconstructions. See Figure 1 – figure supplement 1 for left hemisphere counterpart.

The extent of each bump was identified based on cortical folding (Figure 2; Figure 2 – figure supplement 1). Convexity (curvature) maps were derived from cortical surface reconstructions of the segmented grey matter of T1 images for each monkey. Along the lateral bank of the STS, each bump was constrained laterally by the gyral crown and medially by the fundus. Along the anterior-to-posterior (AP) axis, each bump was constrained to all adjacent cortex surrounding the peak convexity (highest point of bump) terminating in the local minimum (troughs) of the convexity map. Local troughs surrounding the bumps were clearest within the sulcus of the STS just lateral to the gyral crown. Points near and along the gyral crown had uniformly large positive convexities (red colors in the convexity map in Figure 2 & Figure 2 - figure supplement 1). In these regions, the boundaries of each bump were identified based on the troughs within the sulcus and the relative low points of the gyral convexity measures. Across monkeys, the mean surface areas for the posterior, middle, and anterior bumps were 83.69mm^2^ (+/− 5.37 SEM), 74.35mm^2^ (+/− 6.81 SEM), and 63.51mm^2^ (+/− 3.32 SEM). The posterior and middle bumps bordered each other in every hemisphere, but there was consistently a gap between the middle and anterior bumps. Additional bumps were identified within the STS. In each monkey, one or two bumps were present in the posterior-most section of the STS just anterior to the lunate sulcus where the STS arches superiorly towards parietal cortex. Another small bump was also present between the middle and anterior bumps in a few monkeys (e.g. Figure 2 – figure supplement 1, monkey M2). These additional bumps were not face selective and were not evaluated further.

**Figure 2.**
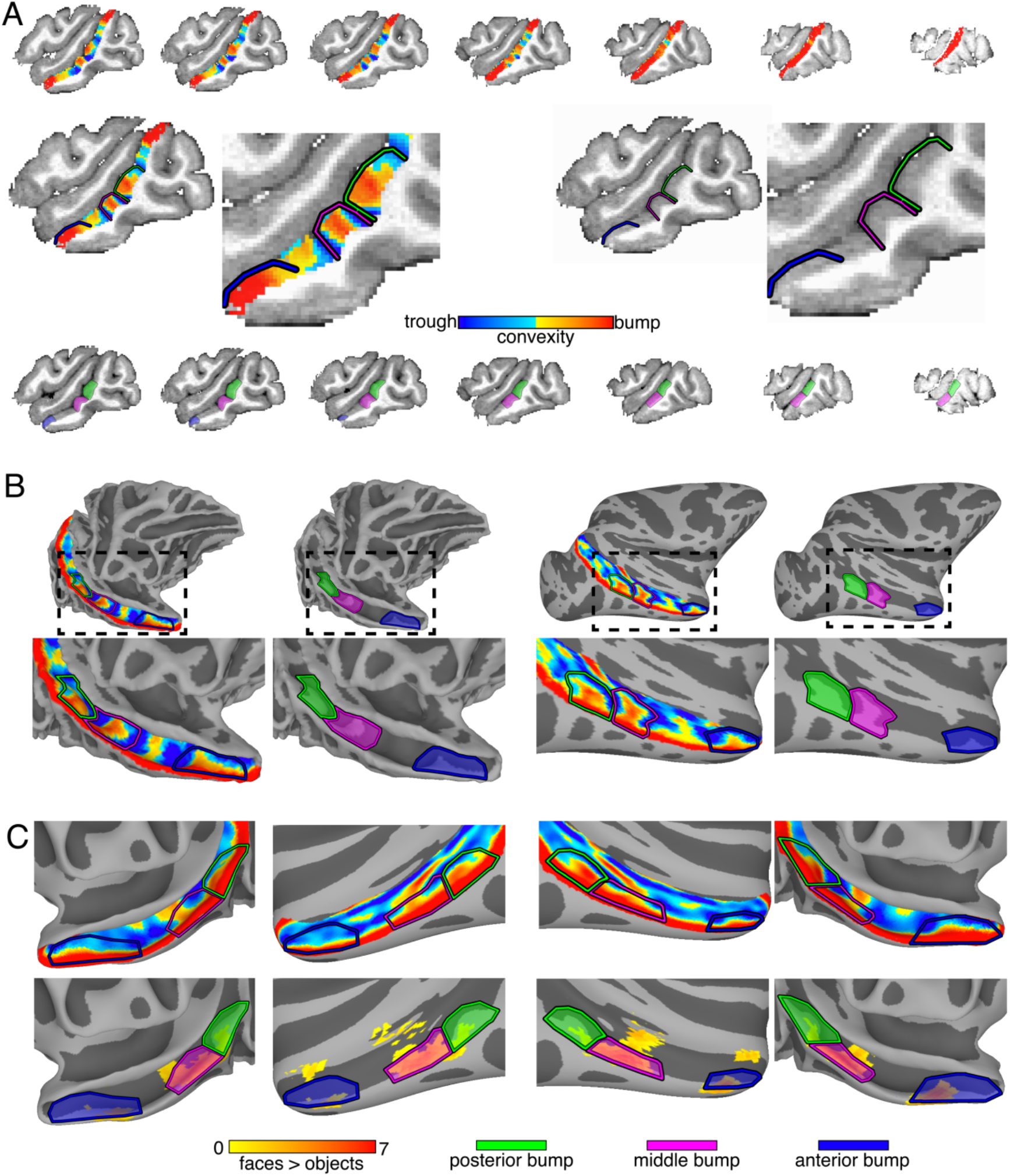
Identification of STS bumps in each monkey. (A) Convexity maps are shown along the lower bank of the STS in seven sagittal slices (1mm spacing) for the right hemisphere of Monkey M2. Enlarged views of single sagittal slices show the bump peaks (red) and troughs (blue). (B) Smoothed white matter and inflated cortical surface reconstructions showing the convexity maps of the STS in the right hemisphere of the same monkey. Posterior (green), middle (pink), and anterior (blue) bumps are shown in both hemispheres. (C, top) Group average convexity maps with outlines of the 3 bumps are shown on the right and left hemisphere surfaces of the NMT template. (C, bottom) Outlines of the three group average bumps are shown overlaid on group average face vs. object activations. The group average map thresholds were set to only show surface nodes where bumps or face selectivity were present in at least 3 monkeys.

Along the posterior-anterior axis, the bumps were localized to similar regions of the STS in each monkey. The posterior bump (Figure 2 – figure supplement 1; green outlined region) was consistently localized to a region of the STS where the inferior occipital sulcus (IOS) intersects the STS and just posterior to where the posterior medial temporal sulcus (PMTS) intersects the STS. The middle bump (pink outlined region) was consistently found anterior to the posterior bump in a region of the STS directly medial and dorsal to the PMTS. The anterior bump (blue outlined region) was found near the anterior tip of the STS directly medial and dorsal to the anterior middle temporal sulcus (AMTS). Across monkeys, the middle bump’s surface area tended to scale with the surface area of PMTS (r = 0.64, *p* < 0.001), even when controlling for total surface area in each hemisphere (partial corr r = 0.61, *p* < 0.001), suggesting a relationship between local morphology of the STS and neighboring sulci. However, such a relationship was unclear for the other two bump-sulcal pairs (rs < 0.28, *ps* > 0.15).

Each STS bump corresponded to an individual face patch. Functional activations were mapped to each monkey’s cortical surfaces (Figure 2 – figure supplement 2) and regions of face selectivity corresponding to previously reported posterior lateral (PL), middle lateral (ML), and anterior lateral (AL) face-selective patches were identified (Tsao et al., 2006; Bell et al., 2009; Pinsk et al., 2009). Each face patch was identified in both hemispheres of all monkeys except for AL in the left hemisphere of Monkey 2 and PL in the right hemisphere of Monkey 4 (Figure 2 – figure supplement 2). The mean surface areas for PL, ML, and AL were 21.37mm^2^ (+/− 3.07 SEM), 57.46mm^2^ (+/− 7.81 SEM), and 44.13mm^2^ (+/− 6.79 SEM). Face-selective regions PL, ML, and AL were localized to the STS bumps. In both hemispheres of each monkey and in the group average data, PL consistently fell on the posterior bump, ML on the middle bump, and AL on the anterior bump (Figure 3). However, in each monkey, face patches were smaller than the bumps, with each face patch covering only a portion of the corresponding bump. Further, some face patches extended laterally in a few individuals. Overall spatial correspondence between face regions and STS bumps was quantified by degree of overlap. For each face patch, a DICE overlap index was calculated with each of the bumps. For PL, ML and AL, the largest overlap was with posterior bump (0.31) the middle bump (0.58), and anterior bump (0.41), respectively (Figure 3, bottom). Together, these data emphasize a correspondence between each lateral face patch and an individual bump in the STS.

**Figure 3.**
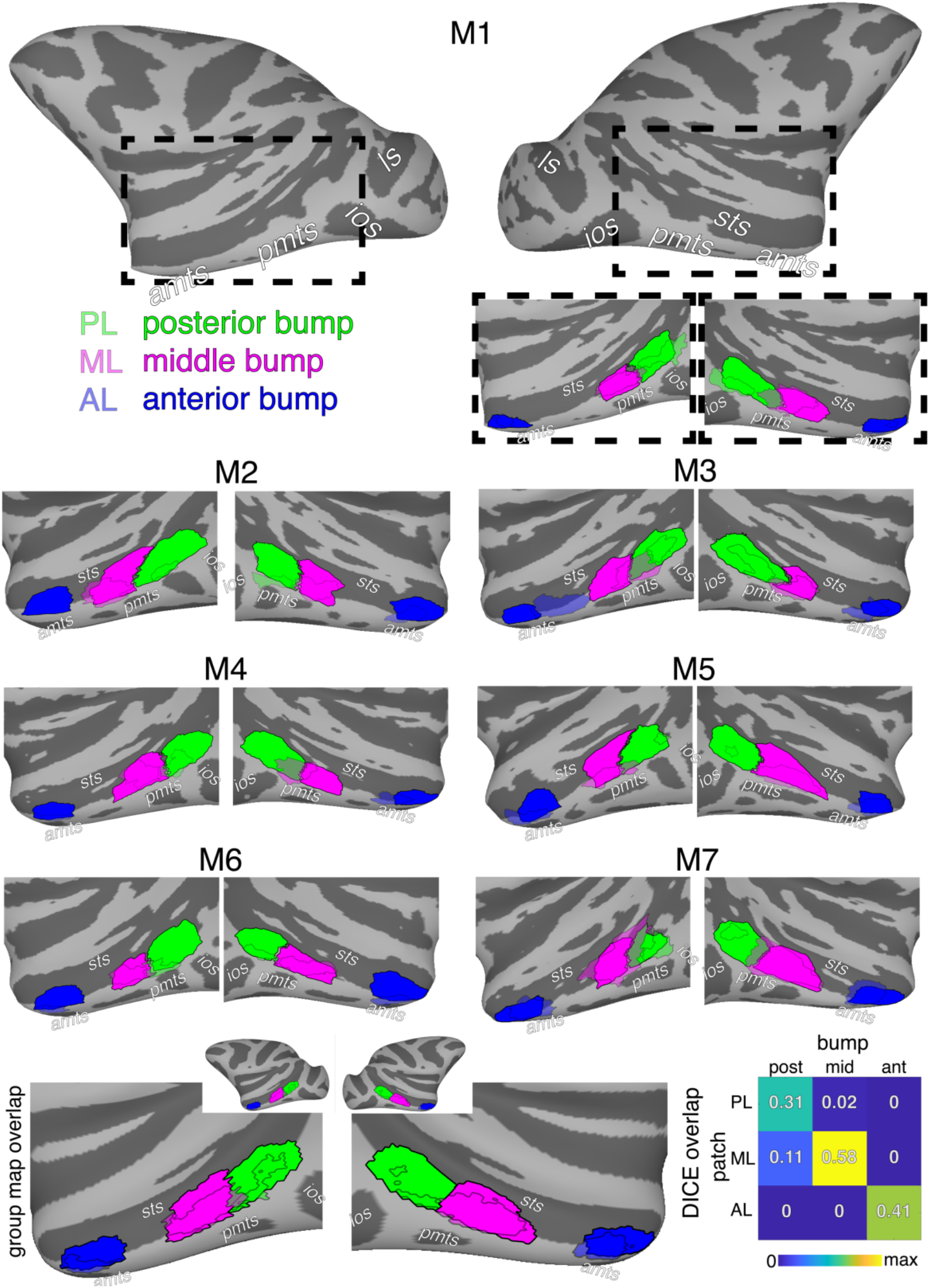
*Overlap between lateral face patches and bumps in the STS. (top) Whole hemisphere inflated cortical surfaces for Monkey 1. Black dashed bounding boxes show the region of STS enlarged for both hemispheres below. (bottom) The spatial extent of STS bumps (opaque colors) and face patches (semi-transparent colors) are shown for both hemispheres in seven normally reared monkeys and* the group average. The group mean DICE overlap indices are shown for all face patch × bump comparisons. *ios* = inferior occipital sulcus, *ls* = lunate sulcus, *sts* = superior temporal sulcus, *pmts* = posterior middle temporal sulcus, *amts* = anterior middle temporal sulcus. Due to the rotation angle, the *amts* is situated underneath the inflated surface in some hemispheres.

Though each bump is in close spatial proximity to a lateral face patch, overlap is not perfect. How does knowing the location of these bumps compare with alternative approaches for localizing face patches? When functional mapping cannot be performed in an individual, the alternative gold standard for localizing a functional region is to use a probabilistic functional atlas where the most probable location of a functional area is calculated from a separate group of individuals. For each monkey, we created probabilistic atlases of each face patch in both hemispheres using the remaining six monkeys. Consistent with a prior study (Janssens et al., 2014), the probabilistic map for each face patch fell within a focal region of the STS. However, there was considerable variability in the spatial overlap of any two monkeys’ face patches. For example, when projected onto the NMT template, the entire right hemisphere ML face patch of Monkey 5 fell anterior to the ML face patch of monkey 4. This lack of spatial correspondence was apparent when looking at face selectivity on the native EPI images. ML falls on the anterior part of the Monkey 5’s middle bump, but on the posterior part of the Monkey 4’s middle bump. Further, few (or no) parts of each probabilistic ROI had 100% overlap across monkeys. On average, the overlap between any individual monkey’s face patch and the corresponding probabilistic atlas (Figure 3 – figure supplement 1, PL, ML, and AL DICE indices = 0.30, 0.46, 0.35, respectively) was worse than the overlap of individual bumps and face patches (Figure 3, bottom). Thus, while probabilistic atlases are useful, similar to the bumps, such approaches are also imperfect at predicting the location of face regions in an individual subject. To directly compare the predictability of bumps with functional atlases, we evaluated the distances between the centroids of each face patch and its corresponding bump / probabilistic ROI. The centroids of face patches were clustered near the centroid of corresponding bumps though most face patch centroids fell closer to the gyral crown of the STS’s lower bank than the bump centroids (Figure 4, top). Consistent with the DICE overlap analysis, the distance along the cortical surface was shortest between each face patch centroid and the centroid of the corresponding bump (PL and posterior bump = 4.28mm; ML and the middle bump = 2.72mm; AL and the anterior bump = 5.42mm). Further, for each face patch, the variance in centroid distances across monkeys was smallest for the corresponding bumps. This indicates that each bump was not just the closest anatomical landmark for its corresponding face patch, but also the most predictive of the face patch’s location. Across monkeys, the overlap was highest and centroid distances were shortest between the ML face patch and middle STS bump indicating that this pair had the strongest structure-function correspondence of the three lateral face regions. The distances between the posterior bump and PL tended to be smaller than the distances between the probabilistic PL and individual monkey’s PL (t(12) = −2.93, *p* = 0.0127). Similarly, the distances between the middle bump and ML tended to be smaller than the distances between the probabilistic ML and individual monkey’s ML (t(13) = −3.16, *p* = 0.0076). There were minimal differences in these distances for AL (t(13) = 0.43, *p* = 0.68). Overall, this suggests that identifying the location of these bumps is comparable to, and in most cases better than, probabilistic atlases.

**Figure 4.**
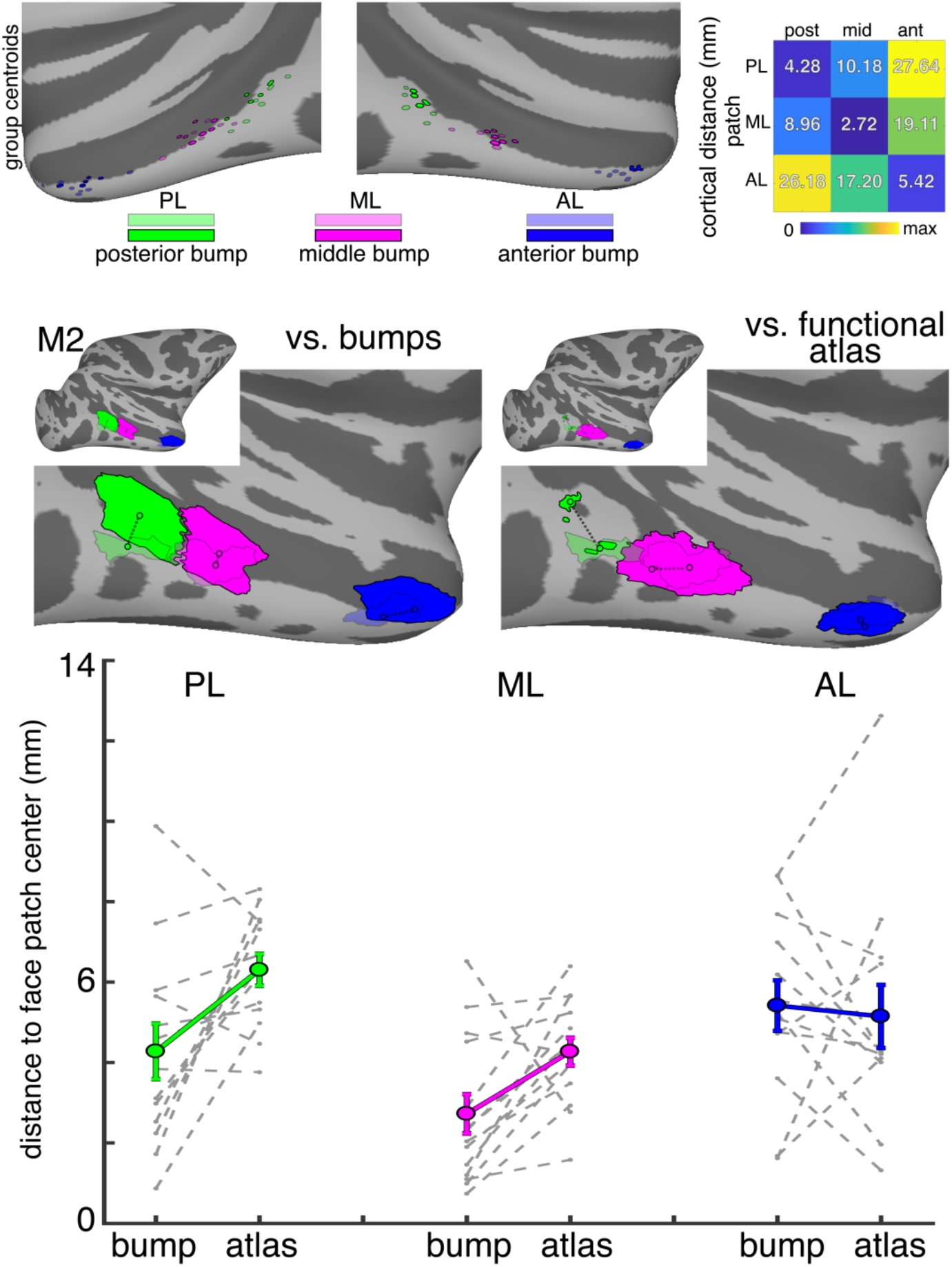
Distances between face regions and STS bumps. (top) The centroids of all monkeys’ face regions and bumps are shown on the NMT template surface. The group mean distances along the cortical surface (in mm) are shown for all face patch × bump comparisons. (middle, left) The distances between the centroids of each face patch and the corresponding bumps are shown for Monkey 2. (middle, right) The distances between the centroids of each face patch and probabilistic functional atlases for each face patch (from a separate group of monkeys) are shown for Monkey 2. (bottom) The distances between bump centroids and face regions and the distances between the probabilistic functional atlas centroids and face regions are shown for individual monkeys (grey dashed lines) and group average (solid colored lines) in PL, ML, and AL.

Each bump contained face-selective neurons. We performed CT imaging to anatomically target where we implanted multi-electrode arrays along the lower bank of the STS in 5 monkeys. Each array was successfully implanted within one of the STS bumps. In each monkey, the vast majority of channels were selectively responsive to faces vs hands, bodies, and inanimate objects (Figure 5). Though we used fMRI in each monkey to verify that face selectivity fell on bumps, we used the bumps as anatomical markers during surgery for our array implantation. The long dimension of each PL and ML array was oriented posterior-to-anterior along the STS and covered the majority of each bump. Further, all arrays targeting face-selective regions have been within bumps and we have yet to implant an array into a bump and not find the majority of channels to be non-face selective. Together, this indicates that the bumps are sufficient for targeting recordings from face-selective neurons.

**Figure 5.**
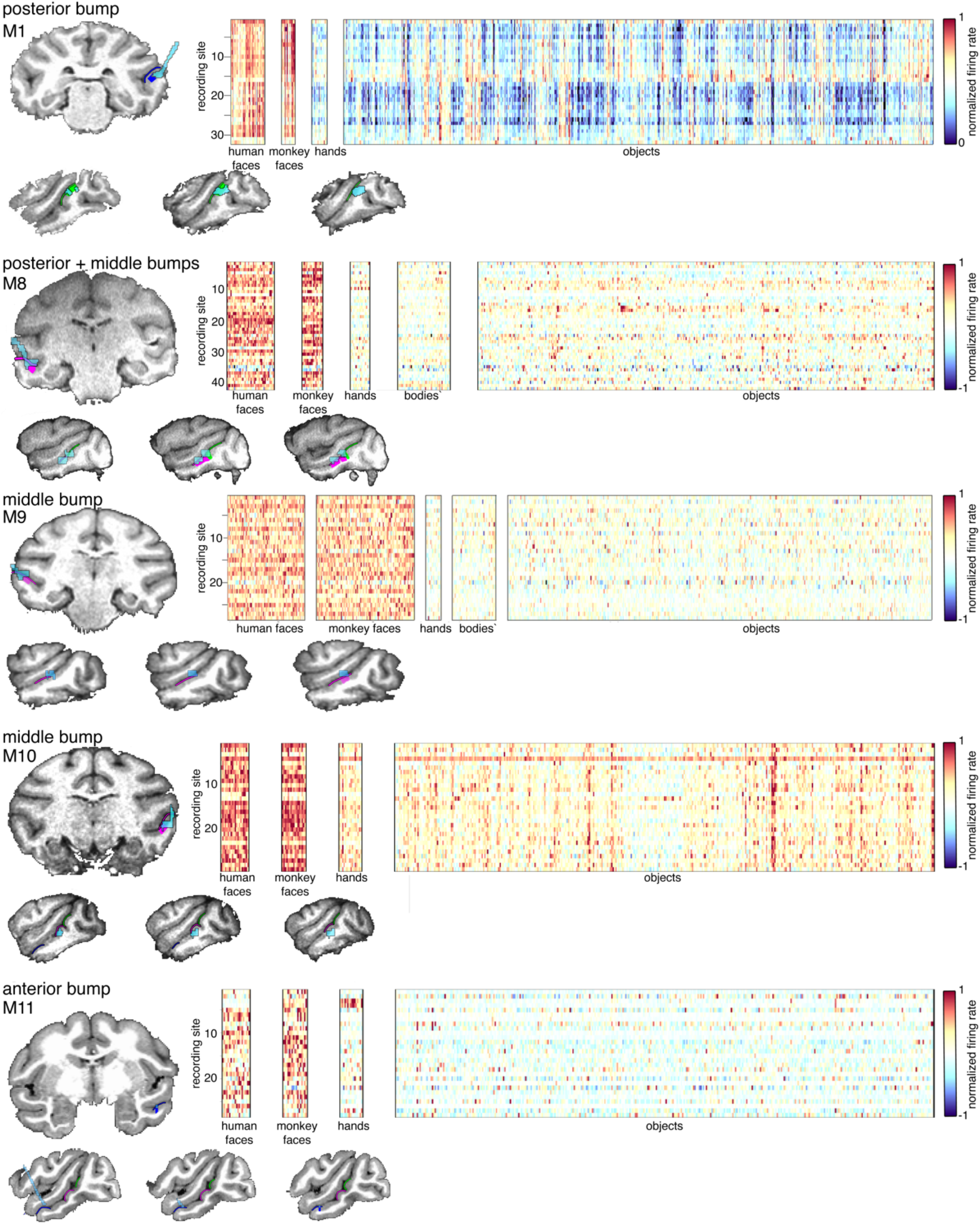
Multi-electrode recordings from STS bumps. Multi-electrode arrays were implanted in the posterior, middle, and anterior bumps of 5 macaque monkeys. In each array, the majority of channels were selectively responsive to human and monkey faces vs. hands, bodies, and inanimate objects.

Though the location of face patches can be predicted by the STS bumps, this anatomical feature of the STS does not necessarily indicate the presence of face selectivity. We collected high resolution T1 anatomical images in 7 monkeys with abnormal early visual experience of faces. Four of these monkeys were raised without seeing faces for the first year of life and did not develop face patches (3 of these monkeys were originally reported in (Arcaro et al., 2017)). Two other monkeys were raised under conditions of general visual form deprivation (not specific to faces) for the first year of life and also did not develop face patches. The seventh monkey was raised in an environment where he had excess exposure of faces to his peripheral visual field during early development (in contrast to the typical foveally-biased visual experience of faces). Though this monkey saw faces and developed face patches, his experience was atypical and serves as a test case of whether the type of face experience affects bump anatomy. In each monkey, the posterior, middle, and anterior bumps were present despite abnormal early experience (Figure 6, Figure 6 - figure supplement 1). Relative to nearby anatomical landmarks such as the IOS, PMTS, and AMTS, the bumps were in similar regions of the STS as compared to the 7 control monkeys reared with normal face experience. Further, the group average bumps from this abnormal monkey group were in good spatial correspondence with the group average bumps from control monkeys (Figure 7). There was a high degree of DICE overlap in the extent of the posterior (0.81), middle (0.73), and anterior (0.77) bumps between the two monkey groups. The distances between centroids of each bump in the abnormal early visual experience group and centroids of the probabilistic face patches were comparable to the centroid distances between bumps and face patches in normally reared individuals (Figure 7, right). The DICE overlap and centroid distances were comparable between the 3 sub-groups of abnormal monkeys: (1) raised without seeing faces, (2) raised with a general visual form deprivation, (3) raised with abnormal constant exposure to faces in the periphery (Figure 7 – figure supplement 1). Together, these data suggest that the STS bumps are present in individuals that lack face selectivity and their macro anatomical organization is similar to that found in monkeys that have face patches.

**Figure 6.**
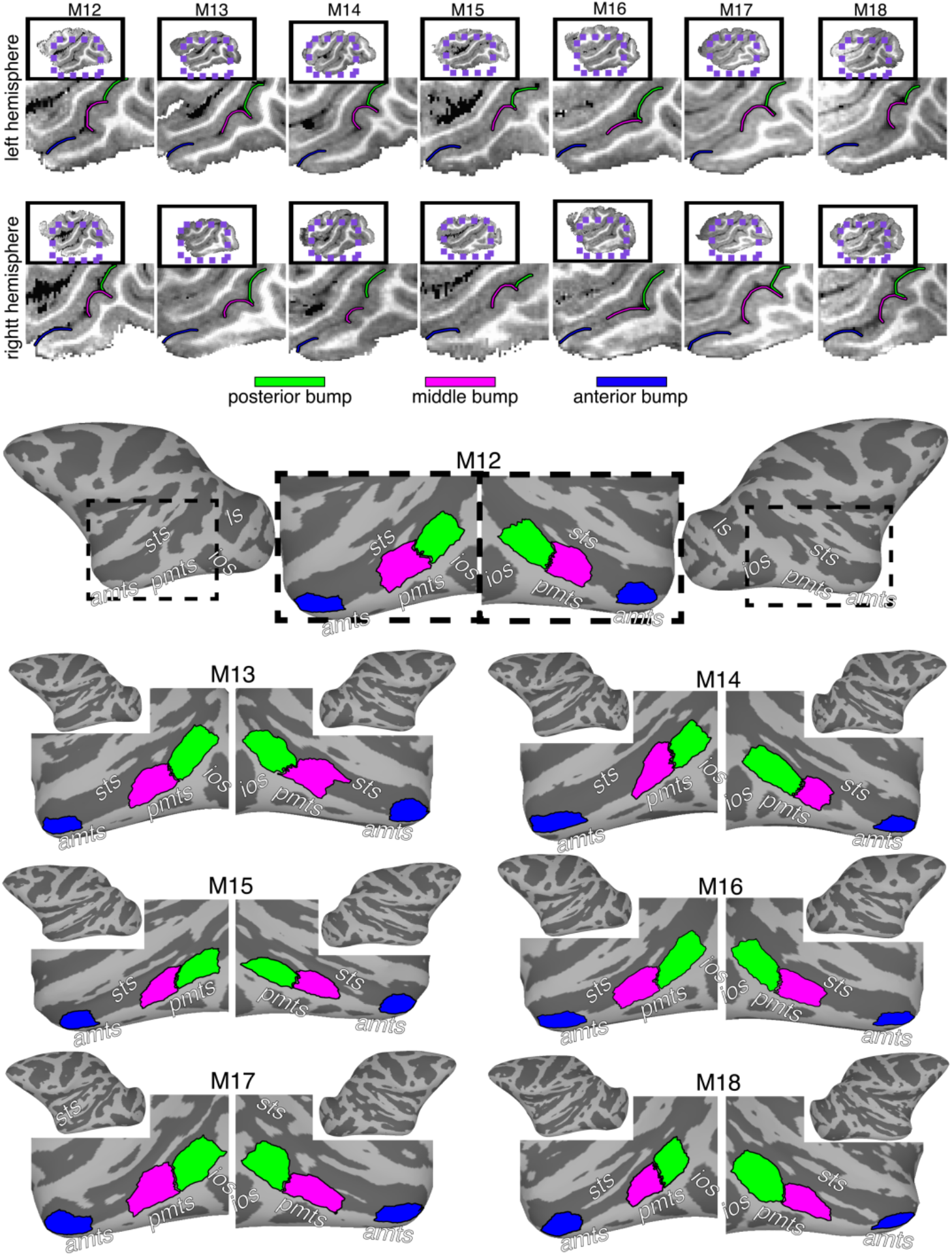
STS bumps are present in monkeys raised with abnormal early visual experience. (top) Single sagittal slices showing bumps in the STS for six monkeys (M12-M17) that lack face-selective regions and a seventh monkey that was raised with an abnormal experience of faces (M18). (bottom) The extent of posterior, middle and anterior bumps is shown on the inflated cortical surfaces for each monkey.

**Figure 7.**
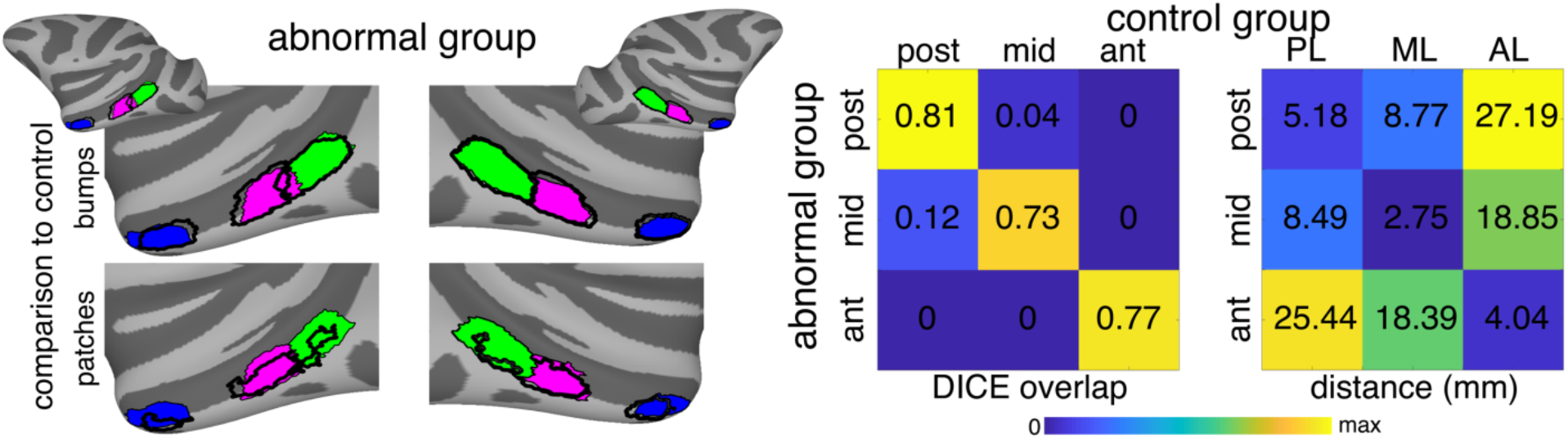
Comparison of bumps between monkeys that have and that lack face-selective regions. (left) The group average location of posterior (green), middle (pink), and anterior (blue) bumps for monkeys raised with abnormal early visual experience are shown on the NMT inflated surfaces. Black outlines show the extent of group average bumps from control monkeys that have face-selective regions. (right) DICE overlap of bumps between the two monkey groups and the group mean cortical distances (in mm) between the centroids of the bumps in the monkeys with abnormal early visual experience and the centroids of probabilistic location of PL, ML, and AL defined from the monkeys reared with typical face experience. See Figure 7 – figure supplement 1 for DICE overlap and centroid distances for the three sub-groups of abnormal monkeys.

STS bumps appear early in development and are evolutionarily preserved across primate species. Though the STS bumps are not clearly formed by gestation day (GD) 110 the sulcus has already begun to bend near where bumps emerge (Figure 8A; pink dashed circle). By GD 135, the posterior, middle, and anterior bumps are all present (Figure 8A, green, pink, and blue arrows). Thus, the bumps emerge prior to seeing faces and well before the development of face patches. The presence of these bumps *in utero* indicates that they form from general principles of cortical folding and cortical expansion. Do these bumps manifest in other primate species? Despite the cortical surface in New World Monkeys being relatively lissencephalic and lacking substantial folds, capuchins have a prominent bump along the middle of their STS anterior to their IOS (Figure 8B). Similar to rhesus macaques, other Old World monkeys including mangabeys and cynomolgus macaques also have clear posterior, middle, and anterior bumps along their STSs. Gibbons and baboons also have similar bumps. Although the cortical surface is much larger in Great Apes and ape brains contain substantially more folds than macaques, gorillas have at least posterior and middle bumps in similar parts of their STS. The topology of the STS is much more complex in orangutans, chimps, and humans and several bumps are apparent. It is unclear which, if any of these bumps, correspond to the three bumps in macaques.

**Figure 8.**
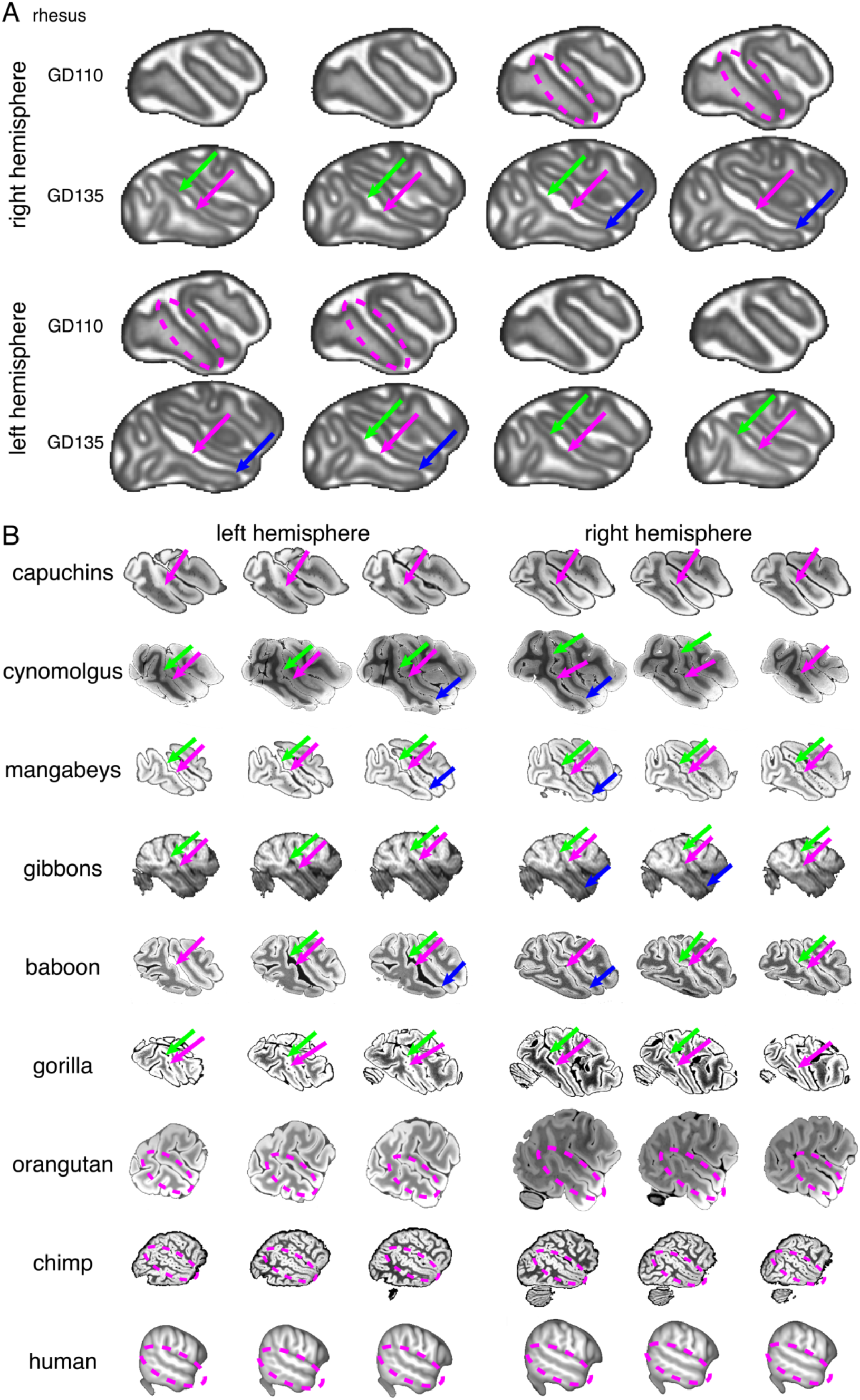
Bump origins. (A) The bump appears early in development. The bumps are not clearly identifiable at gestation day (GD)110 but are present by GD135. Green, pink, and blue arrows indicate posterior, middle, and anterior regions of the STS with bumps. MRI images part of the ONPRC Fetal Macaque Brain Atlas (B) Bumps along the STS are present in several primate species spanning New World Monkeys, Old World Monkeys, gibbons, and baboons. The gorilla STS has bumps similar to macaques, but all other Great Apes have a much more complicated folding structure with several bumps along the STS. All MRI images except for the gibbons come from T2W scans. Gibbons are from T1 images. Dashed circle shows the STS in Great Apes. Gibbon MRI taken from the National Chimpanzee Brain Database (https://braincatalogue.org/). Human MRI taken from the T1 group average from the HCP 1200 Young Adult dataset (https://www.humanconnectome.org/study/hcp-young-adult). All other MRIs taken from the Brain Catalogue (https://braincatalogue.org/).

## Discussion

Our results show a correspondence between the location of face patches and focal protrusions, which we refer to as bumps, along the surface of the STS. In each monkey we identified three bumps. These posterior, middle, and anterior bumps correspond to the fMRI-defined PL, ML, and AL face patches, respectively. Previous research has shown that these face patches fall within similar regions of the STS across individuals (Janssens et al., 2014), but correspondence with specific anatomical landmarks had not been reported. Our analyses indicate that using the bumps to localize face patches in individuals is superior to using a functionally defined probabilistic atlas derived from a separate group of monkeys. Further, recordings from implanted arrays confirmed the prevalence of face-selective neurons within each bump. Together, we demonstrate a viable way to target face patches in the macaque brain based solely on anatomy.

The precise location where each face patch fell on a given bump varied across individuals. Face patches typically fell on the lateral half of the bumps though face-selective activity was found medial to the lateral patches (Figure 2) and likely corresponds to the middle fundus (MF) and anterior fundus (AF) face patches. More notably, face patches varied in their localization along the AP axis of the bumps. This variability was evident in the EPI images (Figure 1 - figure supplements 2 & 3) and thus cannot be attributed to misalignment with the high-resolution anatomical images. It is also unlikely that any differences in vasculature across individuals could account for such variability since the peak of face-selective fMRI activity corresponds to the highest concentration of face-selective neurons (Bell et al., 2009). Instead, this likely reflects true variability in the precise anatomical location of face patches across individuals. Given that these face patches are dependent on face experience (Arcaro et al., 2017) and adhere to a retinotopic proto-organization across IT (Arcaro and Livingstone, 2017), such variability may simply reflect idiosyncrasies of each monkey’s visual experience with faces and how it mapped onto an intrinsic functional architecture. Indeed, a recent study in humans showed a correspondence between face viewing behavior and the receptive field properties of face-selective regions (Gomez et al., 2018). It remains to be seen whether STS bumps in macaques undergo morphological changes across postnatal development and if such changes have any relationship to functional maturation. Thus, it is plausible to have a consistent structure-function correspondence across individuals where the presence and precise localization of function is dependent on how experience interacts with an intrinsic architecture.

These findings add to a growing body of literature demonstrating a surprising degree of structure-function correspondence between cortical topology and the functional organization of higher-order visual cortex (Hasnain et al., 2001; Grill-Spector and Weiner, 2014; Witthoft et al., 2014). Recent research has shown that local, sulcal features are useful for predicting the location of functional domains in human ventral temporal cortex (Weiner et al., 2014; Weiner and Yeatman, 2020). Specifically, the anterior tip of the mid-fusiform sulcus is predictive of the location of face-selective area FFA-2. From this work, one might assume that macaques also would have an anatomical landmark for face patches. However, the sulcal and gyral topology of face-selective cortex differs substantially between humans and macaques. Macaques lack a fusiform cortex and the putative homologous face-selective patches are found in an entirely different sulcus, the STS (Tsao et al., 2008). Thus, it is unclear whether and how such correspondences would hold across these two species. Further, here we find that face patches are anchored to convex bumps, not sulci. Thus, face patches map to particular anatomical features in both primate species, but the specific landmark and even the general property of cortical folding (convex vs. concave) may differ. This is particularly notable since sulcal and gyral regions are hypothesized to broadly differ in their myeloarchitecture (Preuss and Goldman-Rakic, 1991) and areal connectivity (Van Essen, 1997) and, by extension, their functional properties (Li et al., 2017). The identification of these structure-function correspondences in multiple species provides new insight for testing the functional correlates of cortical folding.

What is the possible mechanistic relationship between the 3 bumps distributed along the STS and face patches? A consistent structure-function relationship could be taken as evidence that the location where face patches develop in IT is pre-determined, and the bumps represent circuitry specialized for processing faces. However, morphologically similar bumps were apparent in monkeys that lacked face patches, demonstrating that STS bumps are not sufficient to produce face selectivity in the absence of face experience. Further, these bumps are present *in utero* prior to the onset of vision, and face patches do not appear until 200 days of age (Livingstone et al., 2017). The prenatal formation of these bumps and the consistency in their anatomical location across individuals are doubtless the result of the same general developmental mechanisms that result in the regularity of cortical folding of the entire brain. These factors must include molecular signaling gradients that can indirectly influence folding by limiting the expandability, stiffness, or thickness of the cortical surface (Bayly et al., 2014) and mechanical pressures such as axonal tension (Van Essen, 1997) or differential growth rates of superficial and deep cortical layers (Richman et al., 1975; Lui et al., 2011; Tallinen et al., 2014). In particular regionally specific growth applied to a species-specific initial geometry may suffice for producing species-specific folds, (Kroenke and Bayly, 2018). If anatomical features are produced by general universal mechanisms, then it is unlikely that any particular anatomical feature is innately determined to carry out some particular function. Instead, we suggest that such structure/ function correlates must be based on similarly universal organizational principles, such as maps and connectivity.

Is there an intermediate-level explanation for the association between face patches and bumps? Like Gallia, inferotemporal cortex is generally divided into 3 parts, anterior, middle, and posterior--AIT, CIT, and PIT. Retinotopic mapping shows several central-visual-field foci along the lower bank of the STS with repeating representations of polar angle indicating the presence of multiple visual maps across these 3 parts of IT (Fig 9; Conway, 2018). Along the STS, there are 3 sets of face patches, adjacent sets of body patches (Bell et al., 2009; Pinsk et al., 2009), and 3 sets of color patches similarly distributed along the STS (Tsao et al., 2008; Pinsk et al., 2009; Lafer-Sousa and Conway, 2013). Thus the 3 bumps reflect some aspect of arealization, with a complement of retinotopy and functionality in each area. Whether folding dictates arealization or the reverse is unknown, though prominent cortical folds demarcate the borders between multiple early visual areas. It has been hypothesized that areal connectivity varies systematically with cortical folding. Axonal connections in cortical folds tend to be relatively short and straight, connecting adjacent gyral walls (Van Essen 1997; Hilgetag & Barbas 2006). The association of face patches, and central visual-field representation, to the development of convex bumps is particularly notable given theories of gyrogenesis (Welker 1990), which propose a correspondence between sulcal folds and cytoarchitectonic boundaries thereby potentially centering functional areas on gyri (Hasnain et al. 2001). The bumps may be related to a relative expansion of central visual field. And/or the neuronal morphology and connectivity within these bumps may support computations particularly well-suited for the processing of high-resolution vision, in general, and faces in particular.

**Figure 9.**
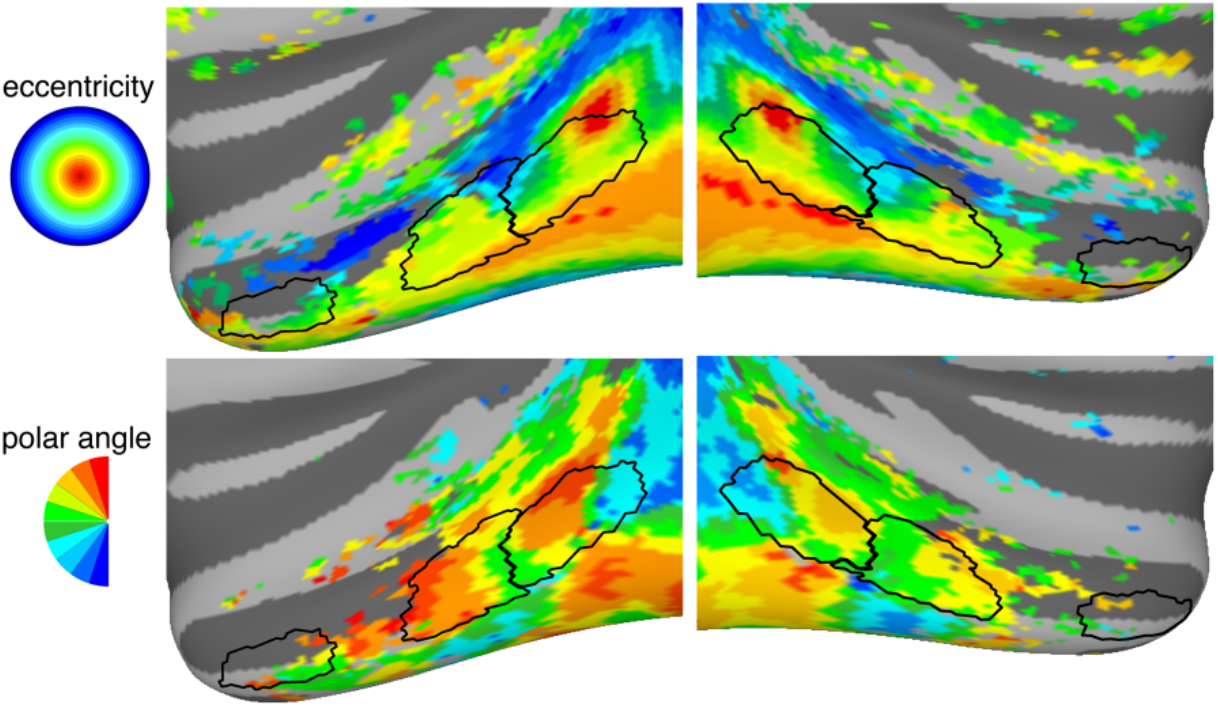
Correspondence between STS bumps and retinotopic organization in IT. (top) Outlines of the STS bumps are shown overlaid on eccentricity maps of visual space that differentiate representations of central (red / yellow) and peripheral (blue) visual space. (bottom) Outlines of the STS bumps are shown overlaid on polar angle maps of visual space that differentiate representations of upper (red), horizontal (green), and lower (blue) meridian representations.

It remains to be seen whether the presence of STS bumps in other animal species indicates the existence or location of face patches. The structure-function topology of visual cortex substantially differs between humans and Old World monkeys. It is generally thought that the face patches within the STS of rhesus macaques correspond functionally to the face patches within the fusiform of humans (Tsao et al., 2008). Though humans also have face-selective regions in their STS (Pinsk et al., 2009), these areas differ in their response properties and are more associated with social and affect features of face processing (Deen et al., 2015; Pitcher et al., 2017). The prominence of these bumps does not fall along strict taxonomy lines, as corresponding bumps were prominent in gorillas but less so in other apes. Identification of such anatomical landmarks may provide insight into evolutionary changes in the functional organization of high-level visual cortex.

## Acknowledgments

We thank K. Weiner for helpful comments on the manuscript. This work was supported by NIH grants RO1 EY 16187 and P30 EY 12196.

## MATERIALS AND METHODS

### Monkeys

Functional and anatomical MRI studies were carried out on 18 Macaca mulatta, 5 female and 13 male. All procedures were approved by the Harvard Medical School Animal Care and Use Committee and conformed with National Institutes of Health guidelines for the humane care and use of laboratory animals. Eleven monkeys (M1-M11) were co-housed with their mothers in a room with other monkeys for the first 4 months, then co-housed with other juveniles, also in a room with other monkeys. Seven of these monkeys (M1-M7) participated in both anatomical and functional neuroimaging experiments. Monkey M1 and the other 4 monkeys (M8-M11) participated in anatomical imaging and electrophysiological recordings from chronically implanted multielectrode arrays. As part of separate experiments, the remaining seven monkeys (M12-M18) were raised with abnormal visual experience of faces. Six monkeys were hand reared by humans for the first year, then were co-housed with other juveniles. Four of the hand-reared monkeys (M12, M13, M16, M17) were raised by laboratory staff wearing welders’ masks that prevented the monkey from seeing the staff member’s face. The only visual experience they had to faces of any kind were during scan experiments, which constituted at most 2 hours per week, with the face exposure being a minor fraction of that time. The other two hand-reared monkeys (M14 & M15) were raised under conditions of visual form deprivation via eye lid suturing for the first year. Lastly one monkey (M18) was raised with his mother in an environment with large posters of faces on the walls such that he had an abnormal, constant exposure to faces in his peripheral visual field. For functional imaging, monkeys were alert, and their heads were immobilized using a foam-padded helmet with a chinstrap that delivered juice. The monkeys were scanned in a primate chair that allowed them to move their bodies and limbs freely, but their heads were restrained in a forward-looking position by the padded helmet. The monkeys were rewarded with juice for maintaining a central fixation within a 2° window. Gaze direction was monitored using an infrared eye tracker (ISCAN, Burlington, MA).

### Electrode implantation

Multi-electrode arrays were implanted within the STS of 5 male Macaca mulatta. Each array was implanted to target face-selective patches. In Monkey 1, a floating microelectrode array (32-channel Microprobes FMA; https://microprobes.com/products/multichannel-arrays/fma) was implanted within middle of the posterior bump of the STS corresponding to his area PL. The internal dimensions of the FMA array was 3.5 × 1.5mm with a spacing of 370-400 microns between electrodes (across the width and length of the array respectively). In Monkey 8, two FMAs were implanted in the left hemisphere – one within the anterior part of the posterior bump corresponding to his PL face patch and a second array within the anterior part of the middle bump corresponding to his ML face patch. In Monkey 9, one FMA was implanted in the left hemisphere centrally within the middle bump corresponding to his ML face patch. In Monkey 10, one FMA was implanted within the anterior part of the middle bump corresponding to his ML patch. In Monkey 11, a 64 channel 12.5 μm NiCr microwire array (McMahon et al., 2014) was implanted centrally within the anterior bump corresponding to his AL face patch. Monkeys were trained to perform a fixation task. Neural recordings were performed on a 64-channel Plexon Omniplex Acquisition System.

### Electrophysiology Display

We used MonkeyLogic to control experimental workflow (https://monkeylogic.nimh.nih.gov). Visual stimuli were displayed on an 19” Dell LCD screen monitor with a 60Hz refresh rate and a 4:3 aspect ratio positioned 54cm in front of each monkey.

### Anatomical imaging

A whole-brain structural volume was acquired while the animals were anesthetized with a combination of Ketamine (4mg/kg) and Dexdomitor (0.02mg/kg). Scans were acquired in each monkey using a 3 T Siemens Skyra, using a 15-channel transmit / receive knee coil. Monkeys were scanned using a magnetization-prepared rapid gradient echo (MPRAGE) sequence; 0.5 × 0.5 × 0.5 resolution; FOV = 128 mm; 256 × 256 matrix; TR = 2700 ms; TE = 3.35 ms; TI = 859 ms; flip angle = 9°). 3 whole-brain T1-weighted anatomical images were collected in each animal.

#### Reconstruction of cortical surfaces

Each animal’s T_1_ images were co-registered to derive an average anatomical volume image for each monkey. Each monkey’s average anatomical volume underwent semi-automated cortical surface reconstruction using FreeSurfer. To ensure high accuracy, skull stripping and white matter masks were first manually segmented by an expert slice-by-slice along coronal, axial, and sagittal planes then passed into FreeSurfer’s autorecon pipeline. Pial and white matter surfaces were inspected to ensure accurate segmentation. If poor segmentations were detected, the white matter mask and control points were edited, and surface reconstruction was rerun until corrected. For several monkeys, FreeSurfer’s autosegmentation had trouble with the calcarine and highly vascularized regions such as the insula. To fix these segmentation errors, average anatomical volumes were manually edited to improve the grey/white matter contrasts and remove surrounding non-brain structures (e.g., sinuses, arachnoid and dura matter).

#### Generation of anatomical feature maps

For each monkey, Freesurfer’s automated algorithm was used to obtain sulcal depth and convexity maps for the pial and white matter surfaces. Sulcal depth (in mm) is measured as the distance between the inflated surface and pial surface (Fischl and Dale, 2000) at each vertex. Convexity maps along the pial and white matter surfaces were obtained using AFNI/SUMA’s automated algorithm (part of SurfaceMetrics). The pial and smooth white matter convexity maps were averaged to produce a mean convexity map.

### Stimuli

Visual stimuli were projected onto a screen at the end of the scanner bore.

#### Static Images

Responses to image categories of faces and inanimate objects were probed. Each scan comprised blocks of each image category; each image subtended 20°×20° of visual angle and was presented for 0.5 seconds; block length was 20 seconds, with 20 seconds of a neutral gray screen between image blocks. Blocks and images were presented in a counterbalanced order. All images were centered on a pink noise background. All images were equated for spatial frequency and luminance using the SHINE toolbox(Willenbockel et al., 2010).

### Scanning

Monkeys were scanned in a 3-T TimTrio scanner with an AC88 gradient insert using 4-channel surface coils (custom made by Azma Maryam at the Martinos Imaging Center). Each scan session consisted of 10 or more functional scans. We used a repetition time (TR) of 2 seconds, echo time (TE) of 13ms, flip angle of 72°, iPAT = 2, 1mm isotropic voxels, matrix size 96×96mm, 67 contiguous sagittal slices. To enhance contrast (Vanduffel et al., 2001; Leite et al., 2002), we injected 12 mg/kg monocrystalline iron oxide nanoparticles (Feraheme, AMAG Parmaceuticals, Cambridge, MA) in the saphenous vein just before scanning.

### General fMRI preprocessing

Functional scan data were analyzed using Analysis of Functional NeuroImages (AFNI; RRID:nif-0000-00259)(Cox, 1996), SUMA(Saad and Reynolds, 2012), Freesurfer (Freesurfer; RRID:nif-0000-00304)(Dale et al., 1999; Fischl et al., 1999), JIP Analysis Toolkit (written by Joseph Mandeville), and MATLAB (Mathworks, RRID:nlx_153890). Each scan session for each monkey was analyzed separately. Using AFNI, all images from each scan session were aligned to a single timepoint for that session, detrended and motion corrected. Data were spatially filtered using a Gaussian filter of 2 mm full-width at half-maximum (FWHM) to increase the signal-to-noise ratio (SNR) while preserving spatial specificity. Each scan was normalized to its mean. Data were registered using a two-step linear then non-linear alignment approach (JIP analysis toolkit) to a high-resolution (0.5mm) anatomical image for each monkey. First, a 12-parameter linear registration was performed between the mean EPI image for a given session and a high-resolution anatomical image. Next, a nonlinear, diffeomorphic registration was conducted. To improve registration accuracy of ventral cortex, we manually drew masks that excluded the cerebellum for both EPI and anatomical volumes prior to registration.

### fMRI Stimulus Category Analysis

A multiple regression analysis (AFNI’s 3dDeconvolve (Cox, 1996)) in the framework of a general linear model was performed on the category experiments for each monkey separately. Each stimulus condition was modeled with a MION-based hemodynamic response function (Leite et al., 2002). Additional regressors that accounted for variance due to baseline shifts between time series, linear drifts, and head motion parameter estimates were also included in the regression model. Due to the time-course normalization, beta coefficients were scaled to reflect percent signal change. Since MION inverts the signal, the sign of beta values were inverted to follow normal fMRI conventions of increased activity are represented by positive values. Brain regions that responded more strongly to monkey faces than familiar objects were identified by contrasting presentation blocks of each of these image categories. Maps of beta coefficients were clustered (>10 adjacent voxels) and threshold at p<0.0001 (FDR-corrected).

### Anatomical analyses

Surface area (mm^2^) was estimated along the pial surface using AFNI’s SurfMeasures. Distances between surface nodes were calculated along the cortical surface using AFNI’s SurfDist.

### Overlap analysis

Spatial correspondence between cortical areas was assessed using the Sorensen-Dice coefficient metric (2|ROI1∩ROI2|/(|ROI1|+|ROI2|)).

### Group Analyses

To directly compare functional data across monkeys, each monkey’s activation maps were aligned to a standard template surface (NMT) using surface-based registration (Freesurfer / SUMA refs). After projecting individual subject data to the template, faces vs objects contrast maps were averaged across monkeys to yield a group average beta map. To visualize group average face selectivity, the data were threshold such that any given surface node needed show significantly greater activity to faces vs objects (*p* < 0.0001, FDR-corrected) in at least 3/7 monkeys. To create group average region-of-interest (ROI) masks for face-selective regions and anatomical bumps, individual monkey masks were projected to the NMT template and averaged. For NMT surface nodes that fell within multiple ROIs across monkeys, the node was assigned to the ROI with the most monkeys. To visualize the group average ROIs, maps were threshold such that any given surface node needed to be within the ROI in 3/7 monkeys. To create probabilistic functional maps of face-selective PL, ML, and AL for each monkey, the face patch ROIs were averaged across all other monkeys.

### Multielectrode array localization

After array implantation, Computed Tomography (CT) scans (0.5 × 0.5 × 1.25mm) were collected. Each monkey’s CT image was spatially aligned to its MPRAGE anatomical image. Because brain/skull contrast is opposite between CT and MPRAGE MRI images, the two volumes were aligned by manually creating a binary brain mask for both CT and MPRAGE images and rigidly aligning the brain masks. The resulting spatial transformation matrix was applied to bring the CT and MPRAGE images into alignment. The locations of the arrays were then compared with the location of STS bumps.

### Multielectrode array analyses

The raw data comprised event (“spike”) times per channel for the entire experimental session. To characterize tuning of each recording site, images of isolated faces, hands, bodies, and objects on a white background were presented within the activating region of all the visually responsive sites. Each image subtended 4° and was presented for 100ms ON and 200ms OFF. Responses were defined as the mean firing rate over 80-250ms after image onset minus the mean firing rate over the first 30ms after image onset. Responses were averaged across image repetitions.

