## Supplemental Figures for "Anatomical correlates of face patches in macaque inferotemporal cortex"

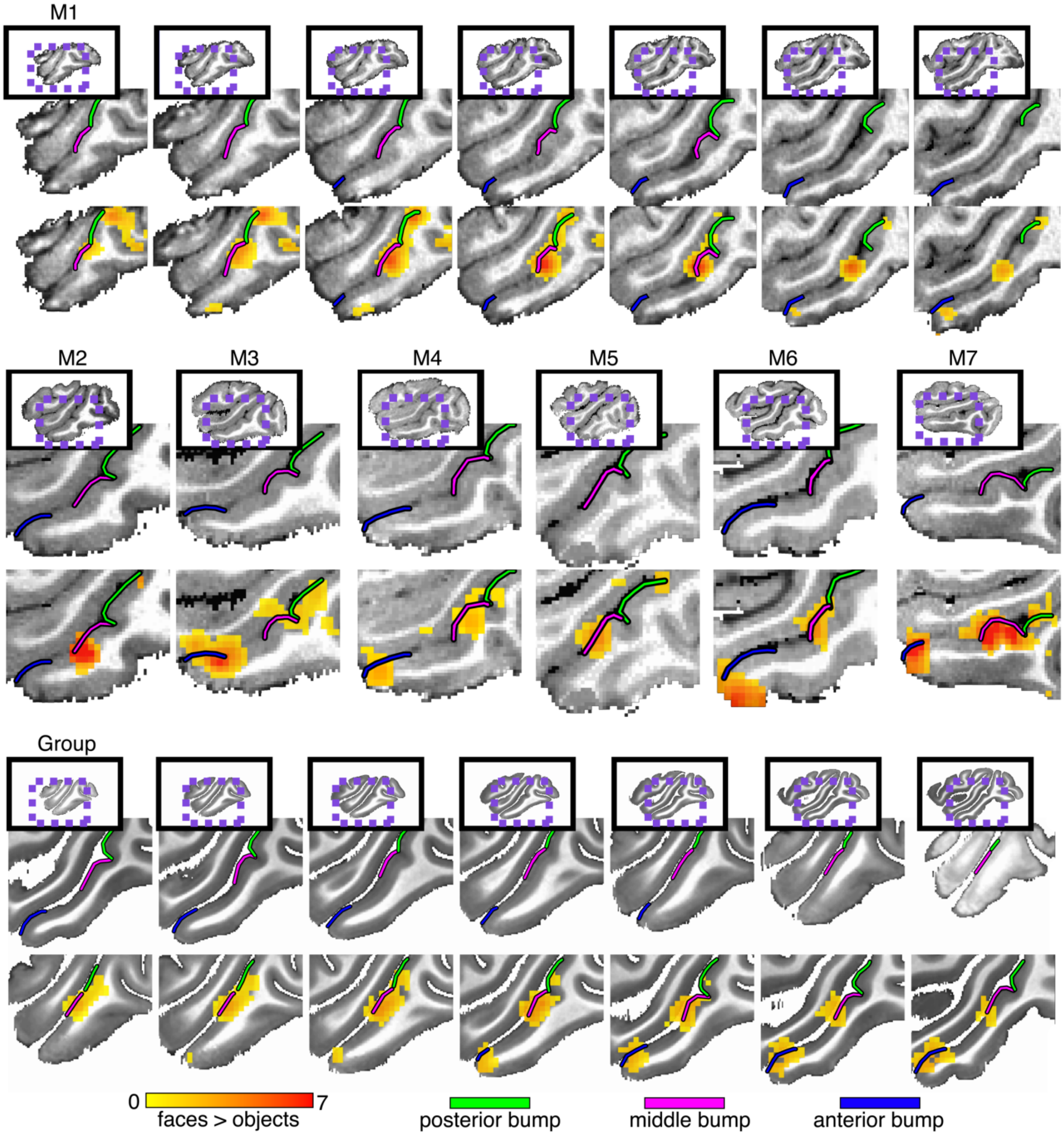


*Figure 1 – figure supplement 1. Anatomical localization of face selectivity in left hemisphere STS. Preferential activity for faces vs. objects was identified along the lower bank of the STS in all 7 monkeys reared with normal visual experience of faces (p < 0.0001, FDR-corrected). (top) Seven sequential sagittal slices at 1mm spacing in the left hemisphere of Monkey 1. (middle) Single sagittal slices showing localization of face-selective activity to bumps in the left hemispheres of the other normally reared monkeys. (bottom) Group average face selectivity falls on anatomical bumps in the STS. Seven sagittal slices at 1mm spacing in the left hemispheres of the NMT template. Group average faces vs. objects (threshold to show only surface nodes where at least 3 monkeys showed significant activity) and convexity maps on cortical surface reconstructions. See Figure 1 for right hemisphere counterpart.*

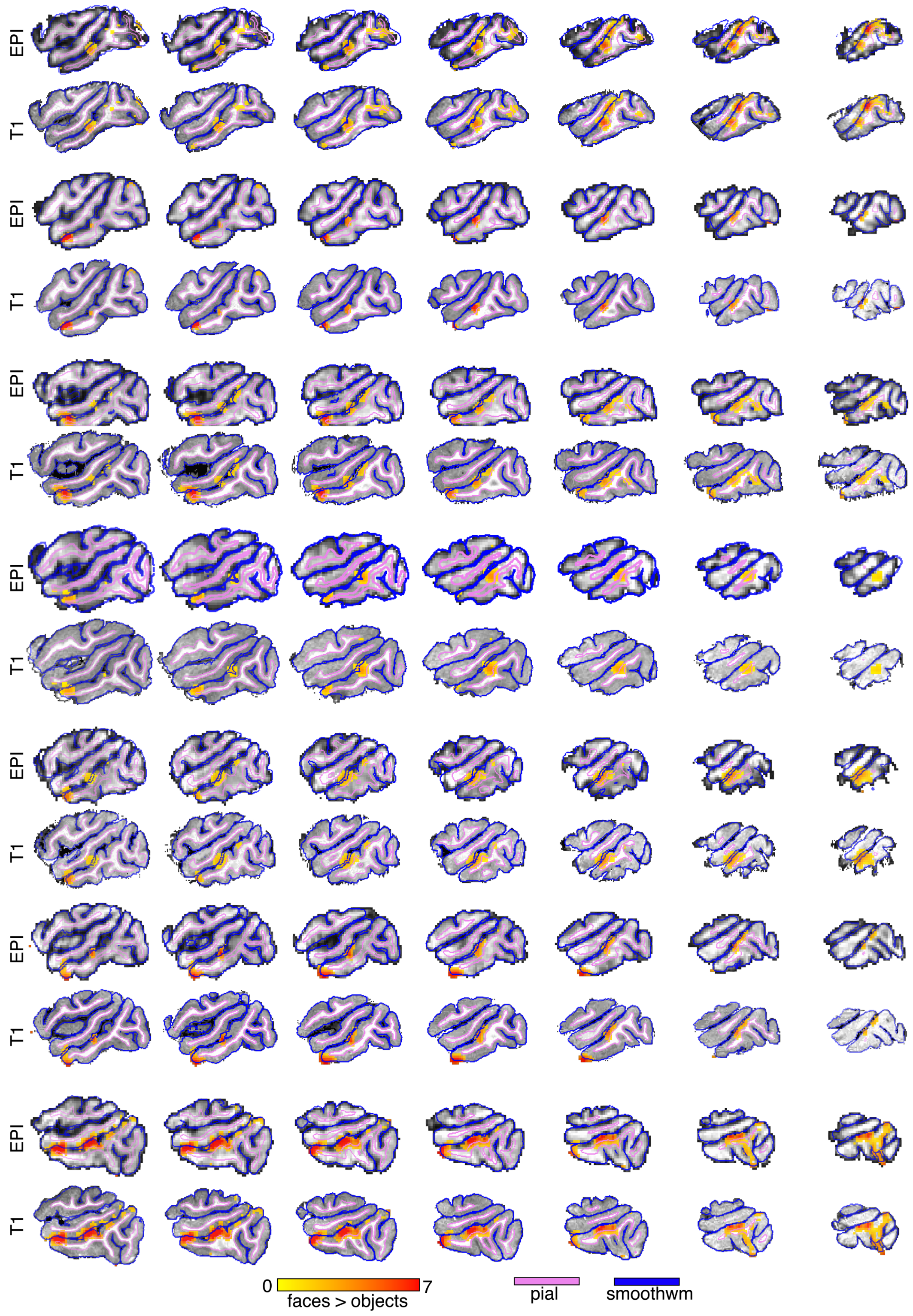


*Figure 1 – figure supplement 2. Face-selective voxels overlaid on EPI and T1 images in right hemisphere of each monkey.*

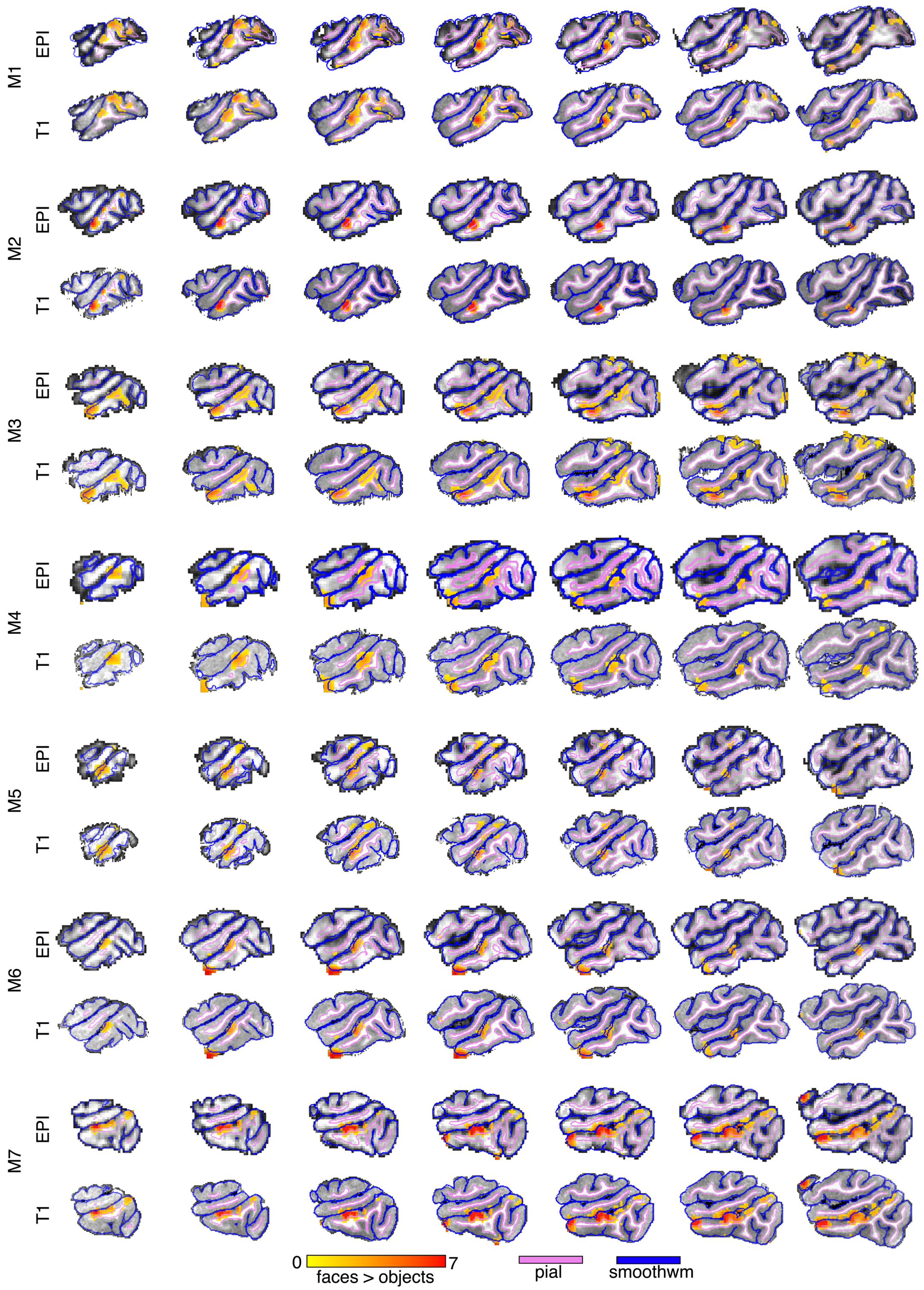


*Figure 1 – figure supplement 3. Face-selective voxels overlaid on EPI and T1 images in left hemisphere of each monkey.*

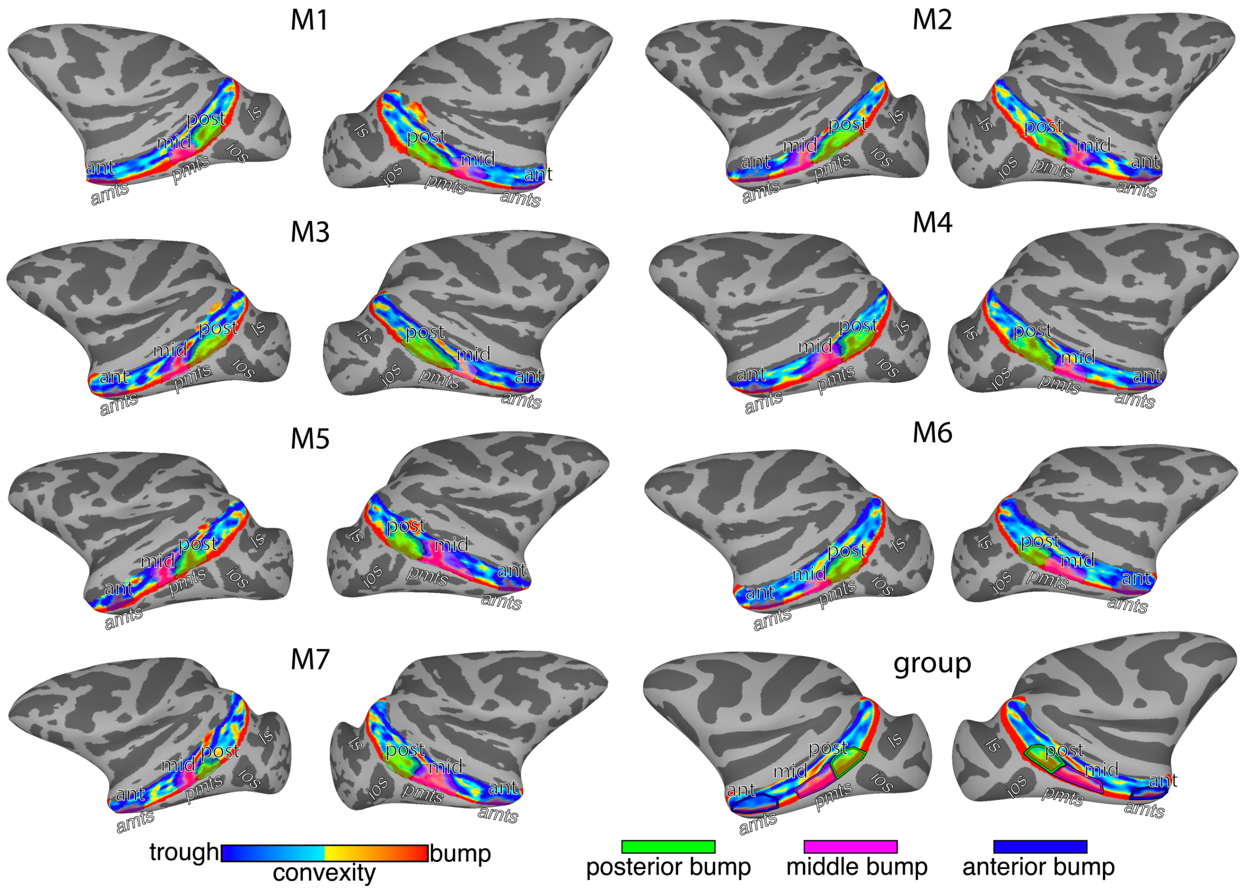


*Figure 2 – figure supplement 1. Convexity maps along the lower bank of the STS in seven monkeys and the group average. Outlined regions on convexity maps correspond to the posterior (green), middle (pink), and anterior (blue) bumps ios* = inferior occipital sulcus, *ls* = lunate sulcus, *sts* = superior temporal sulcus, *pmts* = posterior middle temporal sulcus, *amts* = anterior middle temporal sulcus.

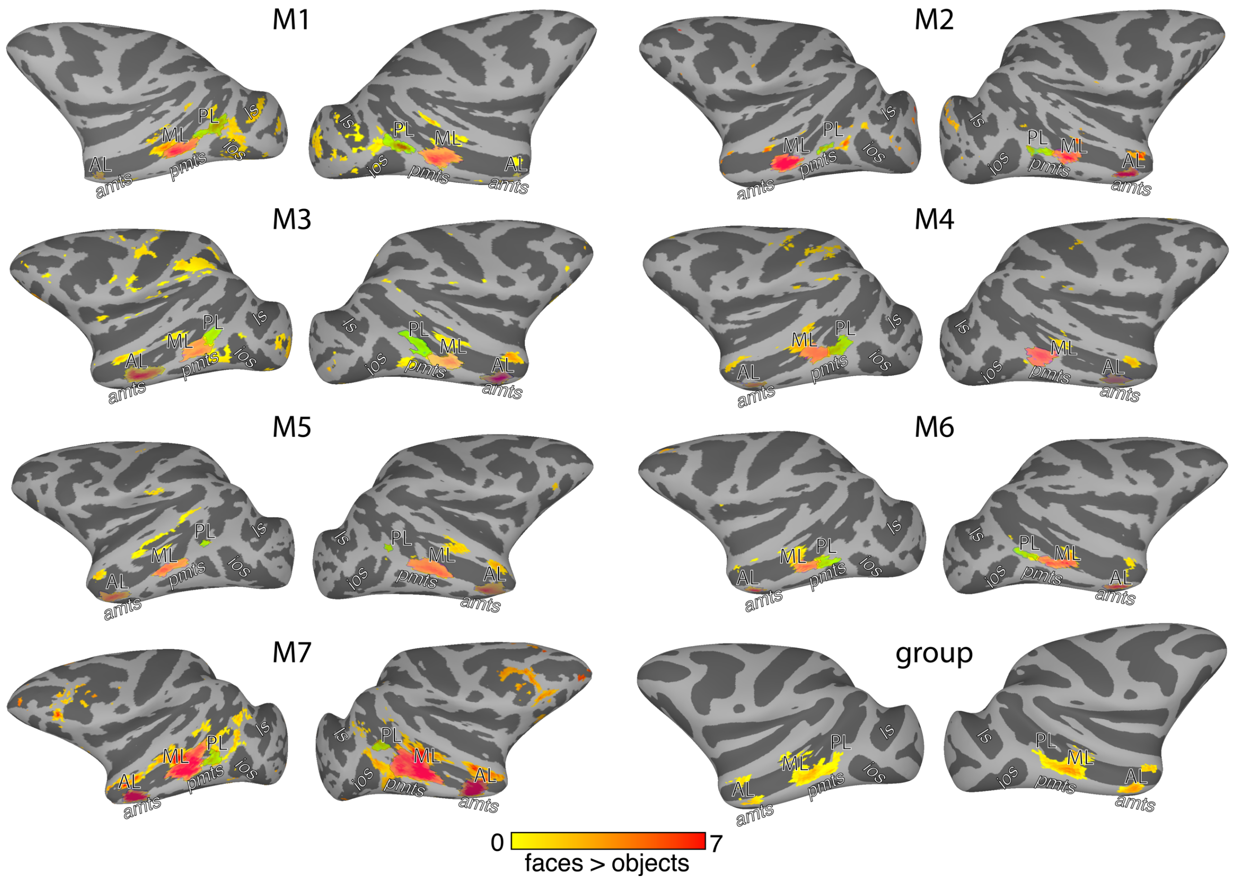


*Figure 2 – figure supplement 2. Face-selective PL, ML, and AL patches along the lower bank of the STS in seven monkeys and the group average. The peaks in face selectivity along the lower bank of the STS correspond to face-selective patches PL, ML, and AL. In each hemisphere, face patches fell on anatomical bumps in the lower bank of the STS. ios* = inferior occipital sulcus, *ls* = lunate sulcus, *sts* = superior temporal sulcus, *pmts* = posterior middle temporal sulcus, *amts* = anterior middle temporal sulcus.

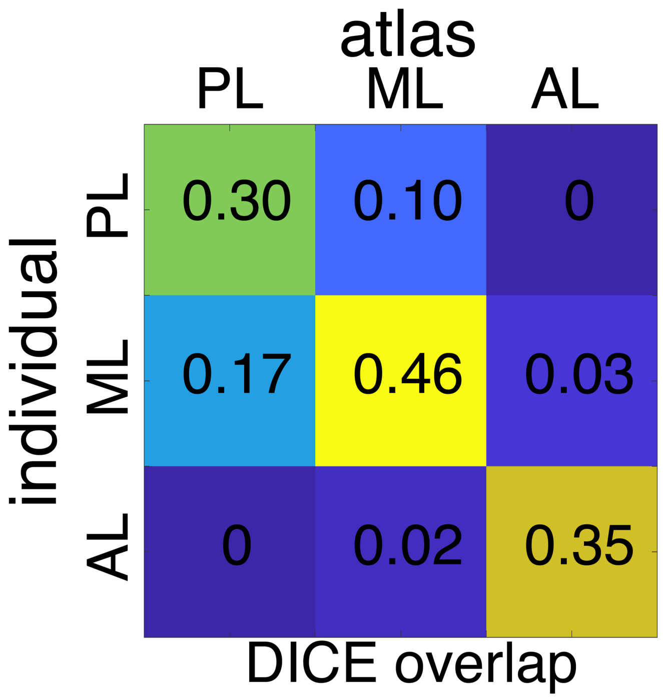


*Figure 3 – figure supplement 1. Group mean DICE overlap between each individual monkey's PL, ML, and AL face patches and the probabilistic atlas.*

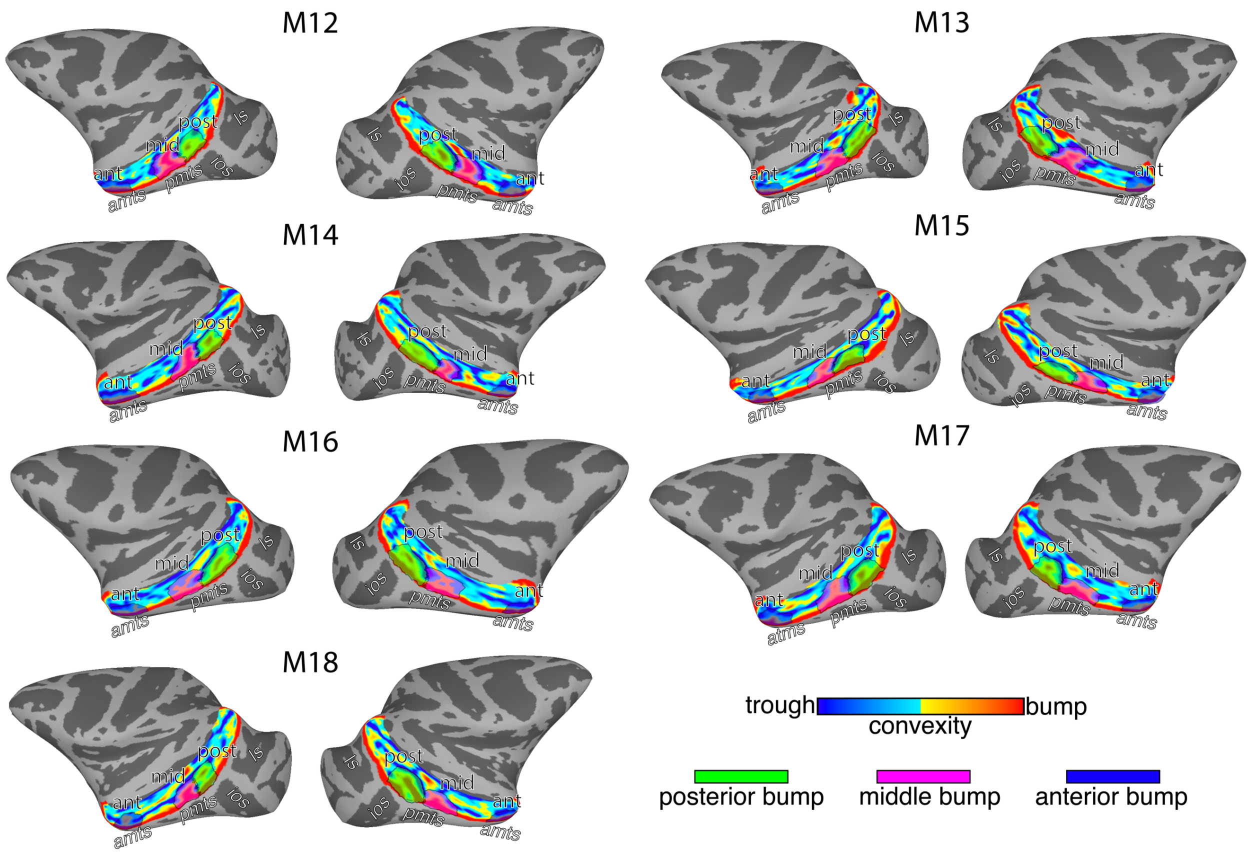


*Figure 6 – figure supplement 1. Convexity maps along the lower bank of the STS in seven monkeys with abnormal early visual experience of faces. Outlined regions on convexity maps correspond to the posterior (green), middle (pink), and anterior (blue) bumps ios* = inferior occipital sulcus, *ls* = lunate sulcus, *sts* = superior temporal sulcus, *pmts* = posterior middle temporal sulcus, *amts* = anterior middle temporal sulcus.

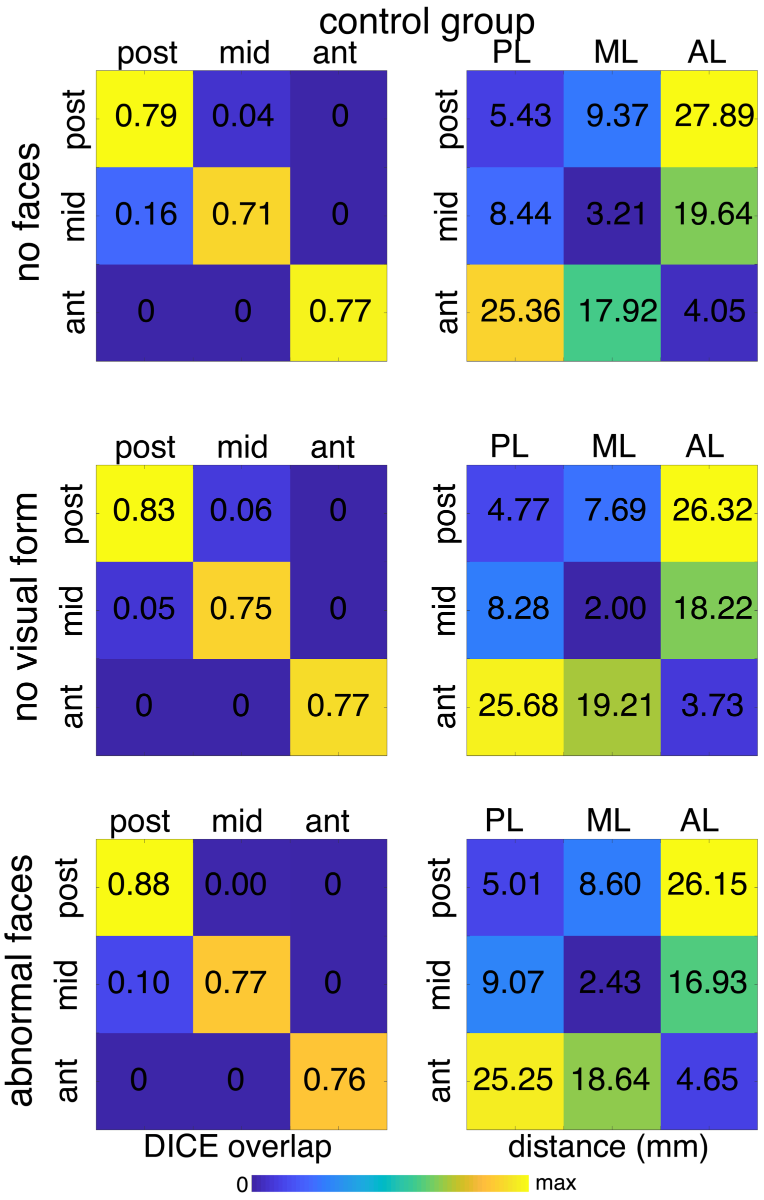


*Figure 7 – figure supplement 1. DICE overlap and centroid distance measures for the three sub-groups of abnormal monkeys. (left column) DICE overlap of bumps between the control and (top) monkeys raised without seeing faces, (middle) monkeys raised under visual form deprivation, (bottom) the monkey raised with abnormal experience of faces in the periphery. (right column) The group mean cortical distances (in mm) between the centroids of the bumps for each abnormal monkey sub-group and the centroids of probabilistic location of PL, ML, and AL defined from the monkeys reared with typical face experience.*
